# Scalable insect cell expression and purification screening applied to CRL4-DCAF substrate receptors

**DOI:** 10.64898/2025.12.04.690205

**Authors:** Angus D. Cowan, Stefan Jaekel, Alessio Ciulli

## Abstract

The ubiquitin-proteasome system is one of the primary mechanisms responsible for degradation of intracellular proteins. Cullin-RING E3 ligases (CRL) are modular, multi-subunit complexes that catalyse ubiquitination of a wide variety of proteins, marking them for degradation by the proteasome. Substrate specificity is conferred by the substrate receptor subunit of the CRL, of which there are hundreds. Targeted protein degradation (TPD) is a drug modality that involves hijacking the activity of CRLs to ubiquitinate non-native neosubstrates via compound-induced ternary complex formation between a substrate receptor and the target. Of the many CRL substrate receptors, the DDB1 and Cul4-associated factor (DCAF) family are of high interest and potential for TPD. To enable characterisation of DCAF proteins and ligand screening campaigns, we have undertaken high-throughput recombinant protein expression screening in insect cells and small-scale plate-based purification of 24 DCAF proteins to identify soluble recombinant protein. Co-expression with the stabilising substrate adaptor DDB1 is required for, or enhances, expression of many DCAFs and provides a folding quality control measure through co-purification with tagged DCAF protein. Of 13 DCAF proteins that had not previously been expressed in the literature, we identify 8 that express well as promising candidates for scale-up. We provide sequence and construct information as a resource for the community. This screening method could be expanded to more DCAF proteins and applied to other CRL substrate receptor families.

## Introduction

Along with transcriptional and translational mechanisms, protein levels in the cell are controlled by protein degradation. The ubiquitin proteasome system (UPS) is one of the primary mechanisms for protein degradation in eukaryotic cells [1, 2]. The UPS involves a set of enzymes that work in concert to mark proteins for degradation by covalently attaching the small protein ubiquitin to lysine residues on the surface of a substrate protein [1]. Ubiquitin is activated in an ATP-dependent manner by an E1 enzyme, to which it becomes covalently bound [3-5]. Ubiquitin is then transferred from the E1 to an E2 enzyme via a transthiolation reaction, forming a thioester intermediate [6-9]. Finally, covalent ligation of the C-terminus of ubiquitin to a lysine residue of a substrate protein is facilitated by an E3 ubiquitin ligase [6, 10, 11]. Polyubiquitination can then proceed where additional ubiquitin molecules are linked to one of several lysine residues on the surface of ubiquitin [12]. Different types of linkages result in different biological outcomes, with K48-linked chains serving as the primary signal that leads to proteasomal degradation [12-14]. Through the UPS, cells can rapidly remove misfolded or damaged proteins and regulate protein levels, both basally and in response to various stimuli [1, 2, 15].

Targeted protein degradation (TPD) involves hijacking the cellular degradation pathways to induce degradation of target proteins [16, 17]. Two classes of well-established degrader molecules, molecular glue degraders (MGDs) and PROteolysis TArgeting Chimeras (PROTACs), function by inducing formation of ternary complex between an E3 ligase, in most cases a Cullin-RING ligase (CRL) [18-21], and a neosubstrate, i.e. a protein not naturally ubiquitinated by the recruited E3 ligase. Formation of the ternary complex leads to ubiquitination of the neosubstrate target protein, which is then recognized by the proteasome and degraded. This modality is already having an impact in the clinic, and many formerly “undruggable” targets are falling to degrader molecules [16, 22-24].

Cullin-RING ligases (CRL) are multi-subunit enzyme complexes that make up the largest family of E3 ligases [18-21]. Over 200 known CRL complexes are classified based on the Cullin scaffolding subunit (Cul1, Cul2, Cul3, Cul4A, Cul4B, Cul5, Cul6, Cul7 and Cul9) [18, 20, 21]. The architecture of an example CRL4 complex is shown in Figure 1. Each Cullin binds an adaptor protein that in turn recruits a substrate receptor subunit [18, 20, 21, 25]. The Cullin bridges these with the E2-recruiting RING protein subunit, facilitating ubiquitin transfer from a ubiquitin-loaded E2 to a substrate protein bound to the substrate receptor [18, 20, 21, 25]. Substrate receptors thereby determine the specific protein or proteins that will be ubiquitinated by a given CRLX^Y^ complex (where X denotes the Cullin scaffold subunit and ^Y^ denotes the substrate receptor subunit). To date, CRL2^VHL^ and CRL4^CRBN^ ligases dominate the MGD and PROTAC landscape, in part due to high quality small-molecule ligands for each substrate receptor, with structurally well-defined binding modes [26, 27].

**Figure 1.**
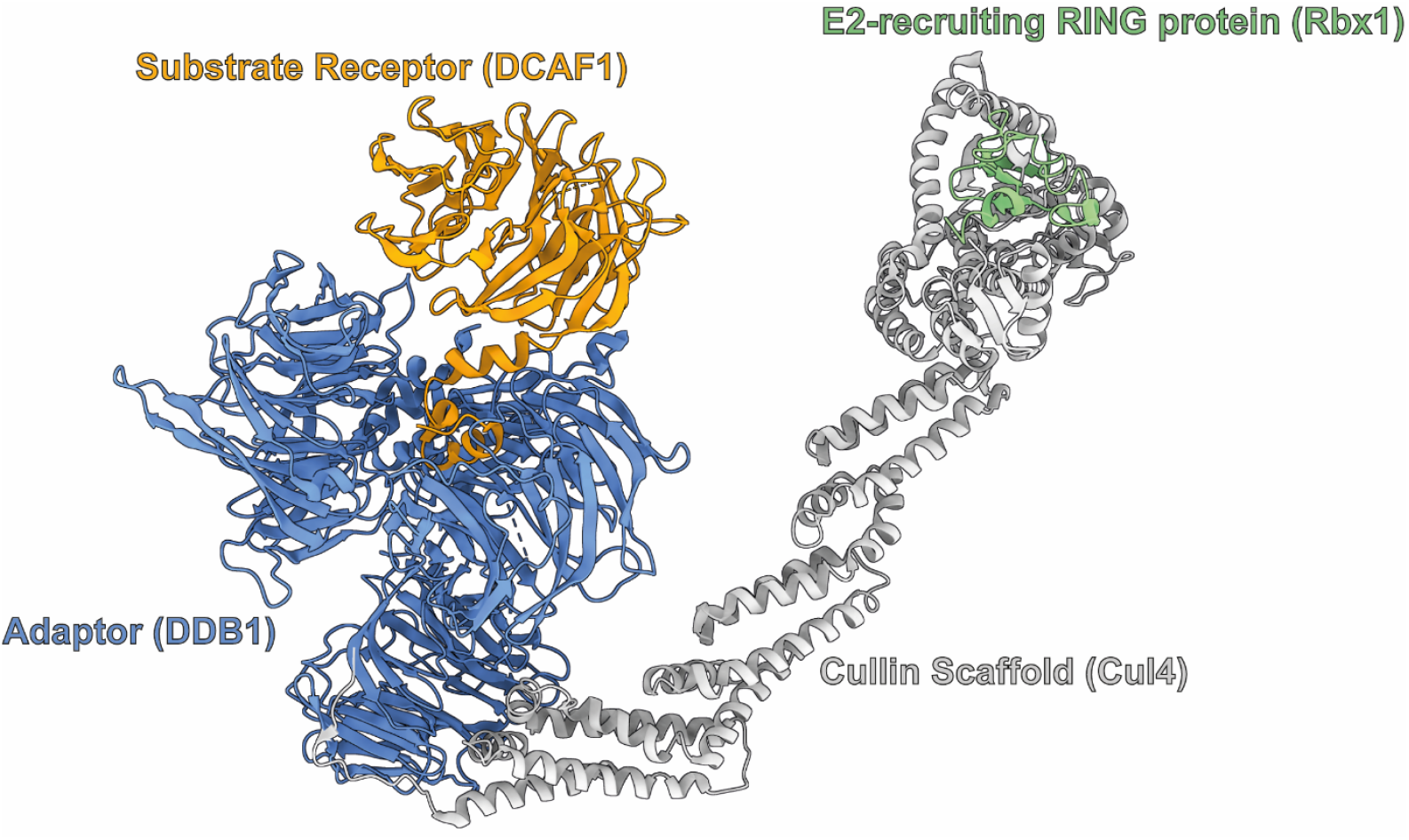
Architecture of an example Cullin-RING ligase (PDB ID: 7OKQ) [28]. The substrate receptor (WDR domain and DDB1-binding helix-loop-helix of DCAF1, orange) that binds substrates for ubiquitination is connected to the Cullin scaffolding subunit (Cullin-4A, grey) via the adaptor protein (DDB1, blue). An E2 ligase loaded with ubiquitin is recruited by the ring protein (Rbx1, green) at the other end of the Cullin protein, positioning it for ubiquitin transfer to the substrate.

Although both ligases have been successfully recruited to degrade a multitude of neosubstrates, the call to expand the toolbox of E3 ligases has been identified as one of the most important challenges in the field of TPD [29, 30]. The choice of E3 ligase affects the degradative potential of a PROTAC or MGD for a given target protein [31-34], which is unsurprising given the importance of forming cooperative and stable ternary complexes between target protein, PROTAC or MGD, and E3 ligase in developing potent degrader molecules [35-38]. Positive cooperativity and stable ternary complexes that facilitate ubiquitin transfer may not be achievable for all potential POIs with the few E3 ligases for which ligands are currently available. Therefore, structural investigation and liganding of other E3 ligases has been identified as an area of critical importance [29]. Beyond ternary complex formation, diversifying the repertoire of E3 ligases available for PROTAC development affords other advantages, including unique temporal, tissue- and organelle-specific expression patterns of different E3 ligases which could enhance specificity and reduce both off- and on-target toxicity [39].

The DDB1 and Cul4-associated factor (DCAF) family of substrate receptors have great potential for development in TPD. DCAF proteins bind the CRL4 adaptor protein DDB1, which can stabilise and enable their recombinant expression, for example expression in insect cells of CRBN and DCAF15 [40-44]. Many DCAF proteins contain a WD40 repeat (WDR) β-propeller domain (Fig 2), a fold with ligandable surfaces and central cavity that makes it an attractive target class for drug development [45]. The ability of DCAF-containing CRL4 complexes to be hijacked for TPD has been extensively demonstrated. Degrader molecules have been discovered and developed for a number of DCAF substrate receptors beyond the most famous example in CRBN, which is extensively recruited as the E3 ligase for MGDs and PROTACs [46-49]. CRL4^DCAF15^ is the recruited ligase for sulfonamide MGDs that degrade RNA splicing factor RBM39 [50, 51]. CRL4^DCAF16^ has been exploited with both covalent and non-covalent degrader molecules, including by covalent PROTACs [52, 53], covalent MGDs [54-57], template-assisted covalent modification [58-60] and intramolecular bivalent glues [61]. Covalent and non-covalent CRL4 ^DCAF11^-recruiting degraders have also been discovered [54, 61-67], and both CRL4^DCAF16^ and CRL4 ^DCAF11^ are frequent hitters in targeted protein degradation screens [68]. Electrophilic PROTACs have been reported for CRL4^DCAF1^ [69], and ligands for CRL4^DCAF1^ and CRL4^DCAF2^ have been discovered and developed into PROTACs [70-74].

**Figure 2.**
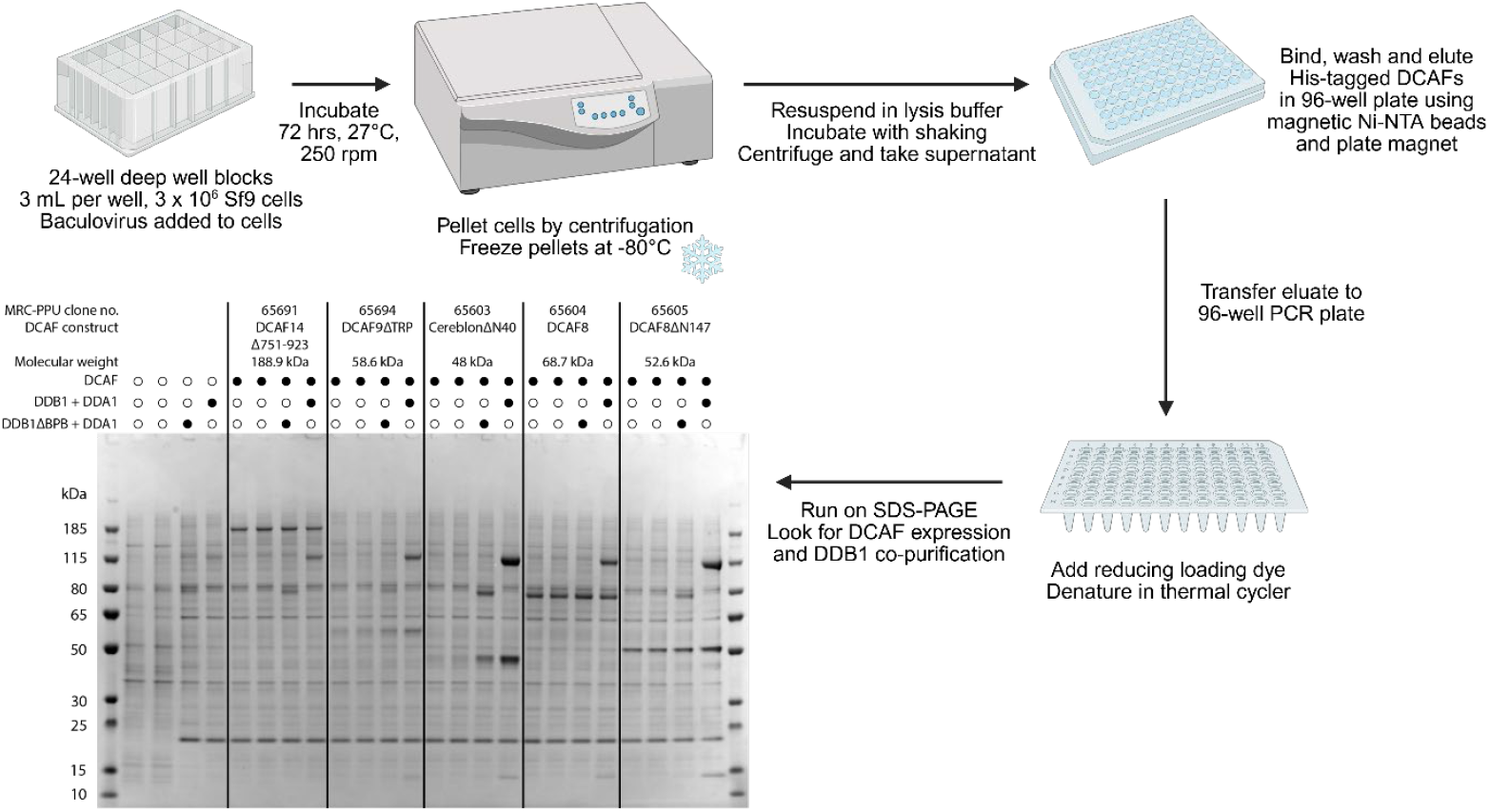
Expression and purification workflow diagram with example gel. Created in BioRender.

Beyond TPD, DCAF proteins are interesting targets in several diseases, including multiple cancer types. DCAF2 (AKA DTL/Cdt2) is implicated in cancer [75] and autoimmune disease [76], DCAF4 (AKA WDR21A) in lung cancer [77], DCAF5 (AKA BCRP2/WDR22) in *SMARCB1*-mutant cancer [78], DCAF7 (AKA HAN11/WDR68), DCAF13 (AKA WDSOF1) in regulating P53 in lung adenocarcinoma [79], and DCAF14 (AKA RepID/PHIP/NDRP/BRWD2) in melanoma, breast and lung cancer [80, 81].

Recombinant expression of proteins is highly enabling for functional characterisation, ligand identification, and drug discovery. Indeed, recombinant DCAF11 and DCAF16 proteins, initially tested and expressed by the methods described here, allowed us to functionally and structurally characterise intramolecular bivalent glue degraders [61]. DCAF proteins have historically proven challenging targets for expression *E. coli*, requiring more complex expression systems such as *Spodoptera frugiperda* 9 (Sf9). Inspired by high-throughput insect cell expression methods from protein production facilities and in the literature [82, 83], we undertook an expression screening campaign of DCAF substrate receptors to accelerate enablement of DCAF E3 ligases for TPD. Our workflow prioritises identification of soluble proteins through rapid small-scale plate-based nickel affinity purification of hexahistidine-tagged DCAF constructs, expressed either on their own or co-expressed with the CRL4 adaptor protein DDB1 and accessory protein DDA1. Co-purification of untagged DDB1-DDA1 by tagged DCAF proteins serves as an indicator of DCAF folding competence and complex formation with their cognate adaptor DDB1.

## Methods

### Construct Design, Cloning and Bacmid Preparation

A total of 54 constructs covering 24 proteins were designed in 2020 (pre-AlphaFold [84]) using secondary structure and disorder prediction software and domain annotations from PredictProtein [85, 86], InterPro [87], and the WDSP database [88-91], or using available structural information for positive controls such as DCAF1, DCAF15 and CRBN [41-44, 92]. A list of constructs and descriptions of their truncations can be found in Table 1. Cloning was performed by the MRC PPU Reagents and Services facility (MRC PPU, School of Life Sciences, University of Dundee, Scotland, mrcppureagents.dundee.ac.uk). The constructs were cloned into pFBDMb, a vector derived from pFastBacDUAL (Invitrogen), under the control of the polyhedrin (polH) promoter with an N-terminal TEV-cleavable hexahistidine tag (MHHHHHHENLYFQGG). DDB1 and DDA1 were cloned into the pFBDMb under the control of the polH and p10 promoters, respectively. DDB1 was cloned either as full-length or DDB1^ΔBPB^, a construct retaining the substrate receptor binding β-propeller domains A and C but lacking the Cullin binding β-propeller B domain (BPB). pFBDMb vectors encoding either DDB1 and DDA1 or DDB1^ΔBPB^ and DDA1 are available from MRC PPU Reagents and Services with code numbers DU70297 and DU70298, respectively. pFBDMb vectors encoding only DDB1 or DDB1^ΔBPB^ without DDA1 (not used in this study) are also available with code numbers DU61229 and DU61230, respectively. Bacmids were generated from donor plasmids in DH10Bac *Escherichia coli* by a standard protocol at the MRC PPU Reagents and Services facility.

**Table 1.** DCAF expression constructs and design rationale.

| Protein | Uniprot ID | Construct | Design rationale |
| --- | --- | --- | --- |
| DCAF1 | Q9Y4B6 | ΔN1020ΔC107 (1021-1400) | Literature published construct, Wu et al. NSMB 2016 |
| DCAF2 | Q9NZJ0 | Full length (1-730) |  |
|  |  | ΔC332 (1-398) | Removes C-terminus beyond blade 7 of WDR domain |
|  |  | Δ188-208ΔC332 (1-187_209-398) | Removes predicted region of disorder between the c to d loop of blade 3 and C-terminus beyond blade 7 of WDR domain |
| DCAF4 | Q8WV16 | Full-length (1-495) |  |
|  |  | ΔN60 (61-495) | Removes predicted region of disorder at N-terminus |
| DCAF5 | Q96JK2 | Full-length (1-942) |  |
|  |  | ΔC580 (1-362) | Removes C-terminus beyond WDR domain |
|  |  | ΔC352 (1-590) | Removes predicted region of disorder at C-terminus |
| DCAF6 | Q58WW2 | Full length (1-860) |  |
|  |  | ΔC21 (1-839) | Removes predicted region of disorder at C-terminus |
|  |  | Δ288-675 (1-287-linker-676-860) | Removes region of disorder between blade 4 of propeller and IQ motif, replacing it with a flexible linker that is commensurate in length to a 27 amino acid IQ motif |
|  |  | Δ288-675ΔC21 (1-287-linker-676-839) | Combination of above truncations |
| DCAF7 | P61962 | Full-length (1-342) |  |
| DCAF8 | Q5TAQ9 | Full length (1-597) |  |
|  |  | ΔN147 (148-597) | Removes predicted region of disorder at N-terminus |
|  |  | ΔC39 (1-558) | Removes predicted region of disorder at C-terminus |
|  |  | ΔN147ΔC39 (148-558) | Removes predicted region of disorder at N- and C-termini |
| DCAF9 | Q8N5D0 | Full-length (1-677) |  |
|  |  | ΔTRP(338-508) (1-337_509-677) | Removes TRP domains |
|  |  | ΔC20 (1-657) | Removes predicted region of disorder at C-terminus |
| DCAF10 | Q5QP82 | Full length (1-559) |  |
|  |  | ΔN119 (120-559) | Removes predicted region of disorder at N-terminus |
|  |  | ΔN119Δ350-396 (120-349_397-559) | Removes predicted regions of disorder at N-terminus and between blades 4 and 5 of predicted WDR domain |
| DCAF11 | Q8TEB1 | Full-length (1-546) |  |
|  |  | ΔN40 (41-546) | Removes predicted region of disorder at N-terminus |
|  |  | ΔC23 (1-523) | Removes predicted region of disorder at C-terminus |
|  |  | ΔN40ΔC23 (41-523) | Removes predicted regions of disorder at N- and C-termini |
| DCAF12 | Q5T6F0 | Full-length (1-453) |  |
| | | ΔN22 (23-453) | Removes predicted region of disorder at N-terminus |
| **DCAF13** | Q9NV06 | Full length (1-445) |  |
| | | ΔC90 (1-355) | Removes Sof1-like domain at C-terminus |
| **DCAF14** | Q8WWQ0 | Full length (1-1821) |  |
| | | ΔC1278 (1-543) | Removes C-terminus beyond WDR domain |
| | | ΔC391 (1-1430) | Removes predicted region of disorder beyond last bromodomain at C-terminus |
| | | Δ751-923 (1-750_924-1821) | Removes predicted region of disorder between WD40 domain and IRS-binding domain |
| | | Δ751-923ΔC391 (1-750_924-1430) | Removes predicted region of disorder between WD40 domain and IRS-binding domain, and predicted region of disorder beyond last bromodomain at C-terminus |
| **DCAF15** | Q66K64 | ΔN29 (30-600) | Removes predicted region of disorder that was not resolved in crystal or cryo-EM structures |
| **DCAF16** | Q9NXF7 | Full length (1-216) |  |
| **DCAF17** | Q5H9S7 | ΔTM (1-185_237-520) | Removes predicted transmembrane regions |
| **RBBP7** | Q16576 | Full length (1-425) |  |
| | | ΔC14 (1-411) | Removes C-terminal region not resolved in crystal structures |
| **DCAF4-like 1** | Q3SXM0 | Full length (1-396) |  |
| **DCAF4-like 2** | Q8NA75 | Full length (1-395) |  |
| | | ΔN20 (21-395) | Removes predicted region of disorder at N-terminus |
| **DCAF8-like 1** | A6NGE4 | Full-length (1-600) |  |
| | | ΔN92 (93-600) | Removes predicted region of disorder at N-terminus |
| | | ΔC38 (1-562) | Removes predicted region of disorder at C-terminus |
| | | ΔN92ΔC38 (93-562) | Removes predicted region of disorder at N- and C-termini |
| **DCAF8-like 2** | P0C7V8 | Full-length (1-631) |  |
| **DCAF12-like 1** | Q5VU92 | Full length (1-463) |  |
| | | ΔN30 (31-463) | Removes predicted region of disorder at N-terminus |
| **DCAF12-like 2** | Q5VW00 | Full length (1-463) |  |
| **Cereblon** | Q96SW2 | ΔN40 (41-442) | Published construct removing predicted region of disorder at N-terminus |

### Baculovirus Generation

A plate-based in-suspension bacmid transfection protocol was adapted and developed from methods described by Roest *et al*. and Scholz & Suppmann for baculovirus generation [82, 83]. Polyethylenimine (PEI), Linear, MW 25000, Transfection Grade (PEI 25K™) (Polysciences) was prepared as per manufacturer instructions at 1 mg/mL and aliquots were stored at -20°C. Baculovirus was generated by transfection of *Spodoptera frugiperda* 9 (Sf9) insect cells with bacmid DNA. For each construct, 6 µg of bacmid DNA was diluted in 200 µL of pre-warmed PBS. PEI 25K (1 mg/mL) was added at a DNA:PEI weight ratio of 1:2 (12 µL PEI per reaction), and the mixtures were mixed gently by inverting the tubes and incubated for ∼20 min at room temperature to allow DNA:PEI complex formation. Meanwhile, Sf9 cells in in log-phase growth were diluted to 0.8 × 10^6^ cells/mL from a maintenance culture. Three millilitres of the cell suspension were dispensed into each well of 24-well Deep-Well Round-Bottom Blocks (Invitrogen). Subsequently, 100 µL of DNA:PEI complex was added per well.

Plates were sealed with AirPore sheets (Qiagen) and incubated at 27 °C with shaking at 250 rpm. At six days post-transfection, cells were assessed microscopically for signs of infection, including cell enlargement and reduced viability. Plates were then centrifuged at 1,500 x *g* in a Beckman Coulter benchtop centrifuge, and the baculovirus-containing supernatants (P0 virus) were harvested, stored at 4 °C, and used within two months.

### Expression

Sf9 cells in log-phase growth were split at a density of 1 x 10^6^ cells/mL one day prior to infection with P0 baculovirus. The following day, 3 mL of the Sf9 cells split again to 1 x 10^6^ cells/mL were added to each well of 24-well deep-well round-bottom blocks (Invitrogen). Each DCAF baculovirus was used both for infection alone and for co-infection with either DDB1-DDA1 or DDB1^ΔBPB^-DDA1 baculovirus. For infection with DCAF alone, 15 µL (0.5% v/v) of DCAF baculovirus was added to the cells. For co-infection, 15 µL (0.5% v/v) of DCAF baculovirus was added together with 15 µL (0.5% v/v) of DDB1-DDA1 or DDB1^ΔBPB^-DDA1 baculovirus, giving a total viral volume of 1% v/v per well. Plates were sealed with AirPore sheets (Qiagen) and incubated at 27 °C with shaking at 250 rpm for 3 days. Infection was confirmed by microscopic inspection of cell morphology. Plates were then centrifuged at 3,273 x *g* for 30 min, the supernatant was removed, and the pellets were frozen in the plates at -80 °C to lyse cells. Pellets were stored in plates at -80 °C until use, usually the following day.

### Plate-based Purification

#### Lysis

For lysis, 16 mL of Binding Buffer 10 (50 mM HEPES, 500 mM NaCl, 10 mM imidazole, 1 mM TCEP, 1% Tween-20, pH 7.5) was prepared per 24-well deep-well block. The buffer was supplemented with one-third of a cOmplete™ protease inhibitor tablet (equivalent to 320 µL of a 1 tablet/mL stock solution), 5 mM MgCl2, and DNase I (grade II from bovine pancreas) (Roche) at twice the manufacturer’s recommended concentration. Defrosted cell pellets were resuspended in 600 µL of the supplemented Binding Buffer 10 per well. Deep-well blocks were shaken at 1,000 rpm for 30 min to ensure complete resuspension and aid lysis. Lysates were clarified by centrifugation of the deep-well block at 3,273 × g for 30 min. Supernatants were transferred to 96-well deep-well blocks (Greiner), and the pellets were discarded.

#### Ni-NTA purification

For purification, 20 µL of Ni-NTA Magnetic Agarose Bead (Qiagen) suspension was dispensed into each well of a flat-bottomed 96-well plate (Qiagen) using a multichannel pipette. Two hundred microlitres of clarified lysate from the 96-well deep-well block was transferred to the corresponding wells, and the plate was shaken at 600 rpm for 1 h to allow binding of hexahistindine-tagged DCAF proteins to the magnetic beads. The plate was then placed on a 96-well magnet (Qiagen Type A) for 2 min, and the supernatant containing unbound protein was carefully removed with a multichannel pipette. After removing the plate from the magnet, 200 µL of Wash Buffer 20 (50 mM HEPES, 500 mM NaCl, 20 mM imidazole, 1 mM TCEP, 0.05% Tween-20, pH 7.5) was added to each well, and the beads were gently resuspended by pipetting. The plate was returned to the magnet for 2 min, and the wash supernatant was removed to a new 96-well microplate. The wash step was repeated once, and the second wash supernatant was discarded. Beads were then resuspended in 50 µL of Elution Buffer 250 (50 mM HEPES, 500 mM NaCl, 250 mM imidazole, 1 mM TCEP, 0.05% Tween-20, pH 7.5), and the plate was shaken for 10 min at 600 rpm to elute bound proteins. Forty-five microlitres of each eluate was transferred to a 96-well PCR plate (non-skirted, Thermo Scientific), mixed with 15 µL of 4x NuPAGE™ LDS Sample Buffer (Invitrogen) supplemented with 200 mM DTT, sealed, and heated to 95 °C for 5 min to denature proteins.

### SDS-PAGE

Twelve microlitres of each boiled sample was analysed by SDS-PAGE on 26-well NuPAGE™ Bis-Tris 4-12% Midi Protein Gels (Invitrogen) in NuPAGE™ MOPS SDS Running Buffer (Invitrogen) with PageRuler™ Plus Prestained Protein Ladder, 10 to 250 kDa (Thermo Scientific).

### Scaled Up Expression and Purification

Expression and purification of DCAF11-DDB1(ΔBPB)-DDA1 and DCAF16-DDB1(ΔBPB)-DDA1 is described in detail in Hsia, Hinterndorfer and Cowan *et al*. [61].

## Results

### Plate-based expression and purification of DCAF proteins

The plate-based trial expression and purification workflow is shown in Figure 2, along with an example gel. Gels for all constructs tested can be found in Supplementary Figures 1-9.

The results of the trial expressions are summarised in Table 1, which reports on the following observables:

- Expression outcome (yes or no)
- Apparent expression level (high, medium, or low)
- Dependence on DDB1 for expression (yes, maybe, or no)
- Enhancement of expression in the presence of DDB1 (yes, maybe, or no)
- Evidence for co-purification with DDB1 and DDA1 (yes, maybe, or no)

Of the 54 constructs tested of 24 DCAF proteins, 48 constructs corresponding to 23 different DCAFs showed detectable expression. Apparent expression levels were estimated qualitatively from SDS-PAGE band intensities relative to the overall protein signal in the same lane and classified as high, medium, or low.

Seven constructs covering 5 DCAFs showed high expression levels: DCAF1, DCAF7, DCAF8, RBBP7, and Cereblon. Of these, human DCAF1, RBBP7, and Cereblon have previously been reported to express in insect cells [41, 93, 94]. Twenty-four constructs covering 12 DCAFs showed medium expression: DCAF5, DCAF8, DCAF9, DCAF11, DCAF12, DCAF13, DCAF14, DCAF16, DCAF8L1, DCAF12L1, and DCAF12L2. Seventeen constructs covering 12 DCAFs exhibited low expression, including DCAF2, DCAF4, DCAF5, DCAF6, DCAF9, DCAF10, DCAF14, DCAF15, DCAF17, DCAF4L 2, DCAF8L1, and DCAF12L1. Finally, DCAF4L1 did not show detectable expression under any condition tested.

Comparison of full-length and truncated constructs indicated that, in most cases, full-length DCAFs expressed at levels comparable to, or higher than, their truncations. Notable exceptions were DCAF2, for which the C-terminal truncation DCAF2ΔC332 expressed while the full-length construct did not, and DCAF8L2, where the N-terminal truncation DCAF8L2ΔN163 showed improved expression relative to the full-length protein.

### DCAF-DDB1 Relationships

DDB1 is the adaptor protein of the CRL4 ubiquitin ligase complex and binds DCAF substrate receptors through various motifs that bind between the DDB1 β-propeller A and C domains, including a conserved helix-loop-helix motif common to WDR domain DCAFs [95, 96] and other unique multi-helical motifs of non-WDR DCAFs CRBN and DCAF16 [40, 58, 61]. Co-expression with DDB1 can stabilise some DCAF proteins which are otherwise insoluble or unstable when expressed in isolation [40-44]. To assess DCAF expression dependence and DDB1 and potential complex formation, DCAF constructs were expressed alone or together with DDB1-DDA1 or DDB1^ΔBPB^-DDA1. DDB1^ΔBPB^ lacks the β-propeller B domain that binds to the Cullin scaffold and is commonly used in structural and biophysical studies of CRL4 substrate receptors [97]. The expression and complex formation relationships between full-length DCAF proteins and DDB1 are summarised in Figure 3.

**Figure 3.**
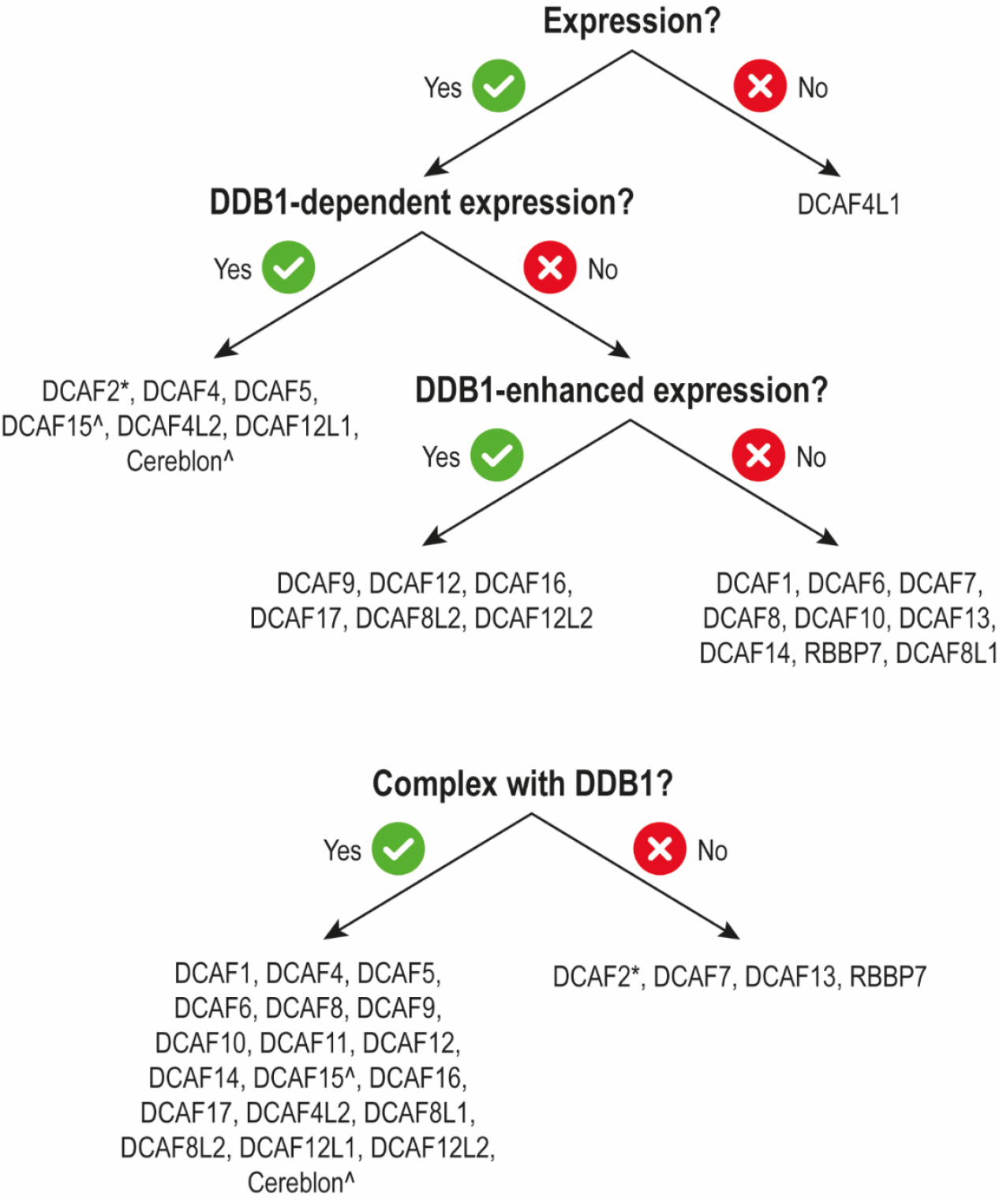
Summary of co-expression and complex formation with DDB1 for full-length DCAF constructs. *Only DCAF2ΔC332 construct expressed and seemed to be dependent on DDB1 virus co-infection but did not form a complex with DDB1. ^DCAF15 and CRBN were literature established N-terminally truncated constructs.

Thirteen constructs showed clear dependence on DDB1 for expression, corresponding to 8 DCAF proteins. All constructs tested for these 8 proteins, including DCAF2, DCAF4, DCAF5, DCAF15, DCAF17, DCAF4L2, DCAF12L1, and CRBN, required DDB1 co-expression to yield detectable levels of DCAF. Expression of 14 constructs was enhanced by co-expression DDB1, including constructs of DCAF6, DCAF8, DCAF9, DCAF12, DCAF16, DCAF17, DCAF8L1, DCAF8L2, and DCAF12L2.

Co-purification analysis identified 40 constructs that retained DDB1 in the eluate, suggesting stable complex formation. These included constructs of DCAF1, DCAF4, DCAF5, DCAF6, DCAF8, DCAF9, DCAF10, DCAF11, DCAF12, DCAF14, DCAF15, DCAF16, DCAF17, DCAF4L2, DCAF8L1, DCAF8L2, DCAF12L1, DCAF12L2, and CRBN.

In contrast, DCAF7, DCAF13, and RBBP7 all showed high expression but did not co-purify with bound DDB1. This lack of interaction could be explained by the absence or disruption of the canonical DDB1-binding helix-loop-helix motif in these proteins. Secondary and tertiary structure predictions of DCAF7 and experimental structures and predictions of RBBP7 lack this motif [84, 94]. DCAF13 also lacks the first helix of the motif, at least when found the human SSU processome complex (PDB 7MQA) [98]. The presence of a proline residue (P30) could disrupt formation of the N-terminal helix of the motif, preventing binding to DDB1. Nevertheless, yeast-two-hybrid screening supports an interaction between DCAF13 and DDB1 [99], and previous studies have observed co-immunoprecipitation of DDB1 with DCAF7, DCAF13 and RBBP7 from human cell lysates [100-104]. Our results suggest DCAF7, DCAF13, and RBBP7 may depend on additional cofactors, post-translational modifications, or alternative interaction surfaces for their recruitment to CRL4 complexes.

DCAF2 was an unusual case, where co-expression with DDB1 did seem to be required for expression of DCAF2ΔC332 (the only DCAF2 construct that expressed) but was not retained in the eluate, possibly because the complex between the two proteins was unstable. A DCAF2 construct consisting of residues 1-458 has been successfully co-expressed and purified with DDB1 in the literature [71].

### Scale up of DCAF expression

Scaling expression of DCAFs tested here should be feasible based on results for DCAF11 and DCAF16 which featured in our work on intramolecular bivalent glues and SMARCA2/4 monovalent MGDs, including structural characterisation by cryo-EM of MGD-induced ternary complexes with DCAF16 [61, 105]. Both DCAF11:DDB1^ΔBPB^:DDA1 and DCAF16:DDB1^ΔBPB^:DDA1 expressed well, with yields up to ∼4.5 and ∼24 mg per litre of High-Five insect cell cultures, respectively. Both DCAF complexes were purified sequentially by cobalt affinity, followed by tag cleavage and reverse cobalt affinity, then anion exchange and size-exclusion chromatography. Example chromatograms and SDS-PAGE gels from the final size-exclusion step are shown in Figure 4.

**Figure 4.**
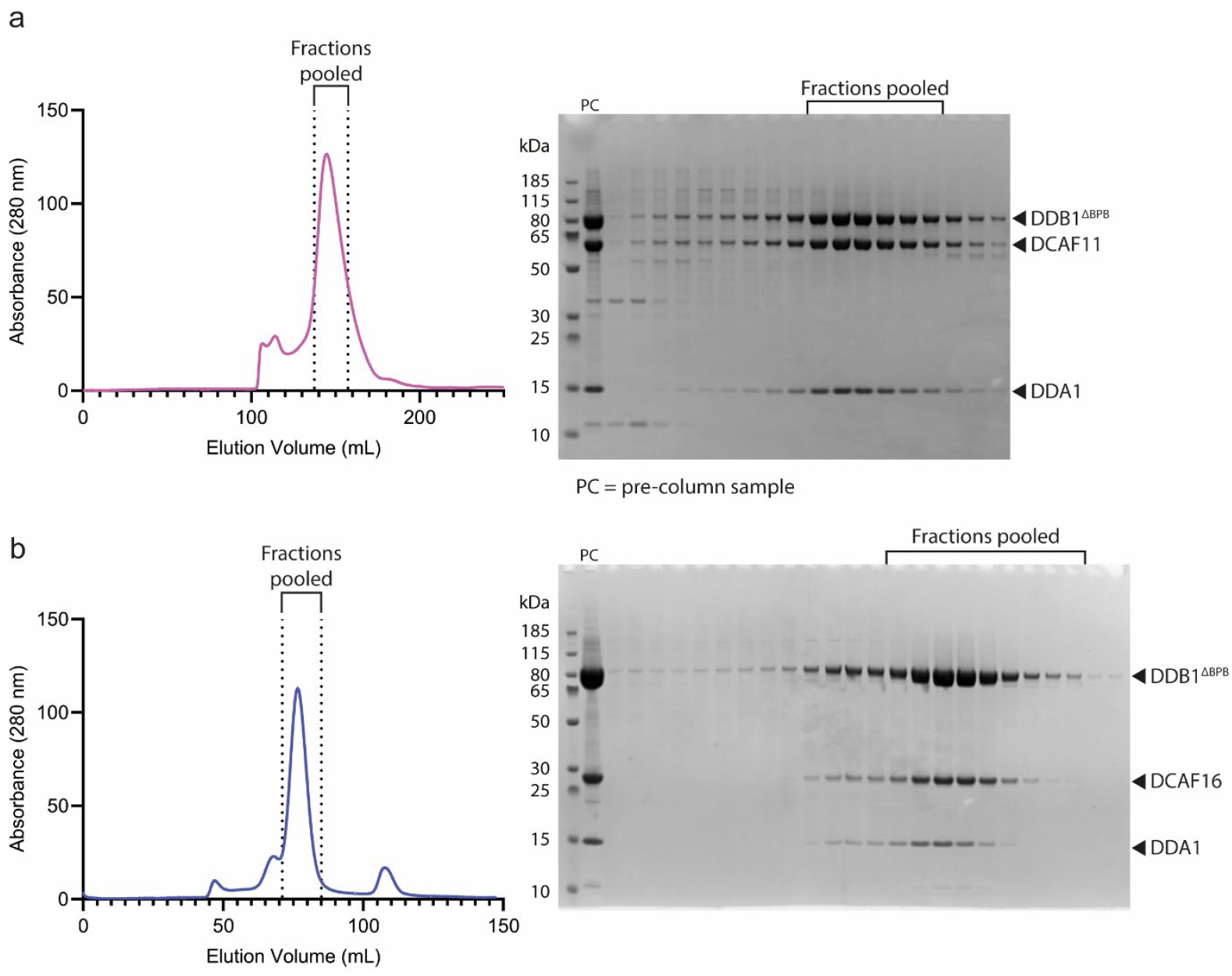
Purification of DCAF11 and DCAF16 complexes by size exclusion chromatography. a) HiLoad 26/600 Superdex 200 pg chromatogram and SDS-PAGE analysis of fractions for purification of DCAF11: DDB1^ΔBPB^:DDA1 complex. B) HiLoad 16/600 Superdex 200 pg chromatogram and SDS-PAGE analysis of fractions for purification of DCAF16: DDB1^ΔBPB^:DDA1 complex. Fractions of complexes pooled for downstream use are labelled. PC = pre-column sample.

## Discussion

Here, we present to our knowledge a first systematic recombinant protein expression screen of a subset of DCAF substrate receptors. Our expression strategy enabled expression of otherwise unstable DCAFs using co-infection with baculovirus encoding the CRL4 substrate adaptor DDB1 and accessory protein DDA1. This workflow also provided internal quality control to assess DCAF folding based on co-purification of DDB1 with tagged DCAF proteins, indicating receptor-adaptor complex formation. Overall, the success rate for expression was high, with 23/24 DCAF proteins showing some level of expression for at least one construct.

The plate-based methods employed from baculovirus generation through to affinity purification are scalable and translatable. The method could be used to screen expression of other E3 ligase substrate receptor families with their cognate adaptors, including ∼75 CRL1 F-box substrate receptors with the adaptor Skp1 and ∼50 CRL2 and CRL5 BC-box substrate receptors with the adaptor ELOB/C. Conceptually, the co-infection/co-expression and co-purification method is applicable in other expression systems (e.g. *E. coli* and mammalian cell expression) and could be applied to any multi-subunit complexes with a core component (analogous the adaptor) and modular interchangeable subunits (analogous to the substrate receptors).

The structures of several DCAF proteins, either full-length or truncated, have been reported in the literature, often in complex with DDB1. They include DDB2 (complexes with DDB1) [95, 106-108], CSA (complexes with DDB1) [106, 109-111], DCAF1 (many structures of WDR with and without DDB1) [28, 69, 70, 73, 74, 92, 93, 112, 113], DCAF2 (complexes with DDB1) [71], DCAF3/AMBRA1 (complexes with DDB1) [114, 115], DCAF5 (complex with DDB1) [78], DCAF12 (complexes with DDB1) [116, 117], DCAF13 (SOF1 in yeast, component of the small ribosomal subunit processome) [98, 118], DCAF14 (bromodomain 2 only) [119, 120], DCAF15 (in complex with DDB1-DDA1 and in some cases RBM39) [42-44, 121], DCAF16 [58, 61, 105], RBBP7 [94], and CRBN (many structures of the thalidomide binding domain, CRBN^midi^ and full-length/near full-length constructs with DDB1) [40, 41, 122, 123]. Recent work from structural genomics consortium focused on evaluating target class ligandability of WDR domain proteins, included DCAF1, DCAF11 and DCAF12 [45]. In agreement with our results, expression for DCAF11 was only reported with co-expression DDB1, as was the case with many of our tested DCAFs.

Many of the DCAF proteins expressed here have not been recombinantly expressed or had structures solved previously, including DCAF4, DCAF6, DCAF7, DCAF8, DCAF9, DCAF10, DCAF11 (recombinantly expressed but no structures), DCAF14 (no structures for full protein/WDR domain), DCAF17, DCAF4L2, DCAF8L1, DCAF8L2, DCAF12L1, DCAF12L2. Of these proteins, we identify DCAF4, DCAF8, DCAF9, DCAF14, DCAF8L1, DCAF8L2, DCAF12L1 and DCAF12L2 as promising for follow up studies, particularly DCAF8 and DCAF14 which expressed well. Liganding of more DCAF proteins such as DCAF8 and DCAF14 holds promise for tissue-specific degradation by DCAF-recruiting PROTACs, as many DCAFs are differentially expressed in different tissues and are highly expressed in the testis [124].

The DCAF constructs tested in this work were designed prior to the advent of AlphaFold [84], although certain unique DCAFs such as DCAF16 had only very poor confidence predictions that proved incorrect. Nevertheless, our work shows almost all the DCAF proteins tested are amenable to insect cell expression, and constructs can likely be further refined using much improved structure prediction from modern AI/ML tools. In addition to the constructs reported here, we also designed and cloned further constructs of DCAF2, DCAF3/AMBRA1, DCAF6, DCAF10, DCAF15, DCAF16, DCAF17, DCAF19, DCAF8L2 and DCAF12L2 (some with the aid of AlphaFold) but did not test their expression. For posterity, these constructs can be found in supplementary table 1 along with all tested constructs, with construct details and clone reference numbers. We anticipate this methodology and shared resource will be of great interest to the growing field of targeted protein degradation and targeting of ubiquitin E3 ligases, and will find utility and value for structural biologist, biochemists and chemical biologists interested in studying the elusive functional roles of DCAFs and finding new small-molecule binding ligands for inhibitor and degrader development.

## Declarations

## Availability of data and materials

All data referred to are presented in the manuscript. All DCAF and DDB1-DDA1 donor plasmids are available from the MRC PPU Reagents and Services facility (MRC PPU, School of Life Sciences, University of Dundee, Scotland, https://mrcppureagents.dundee.ac.uk).

## Funding

This work was funded by the pharmaceutical companies supporting the Division of Signal Transduction and Therapy (Boehringer Ingelheim, GlaxoSmithKline, Merck KGaA) as sponsored research funding to A.C. Funding in the Ciulli Lab is also gratefully acknowledged from the Innovative Medicines Initiative 2 (IMI2) Joint Undertaking under grant agreement no. 875510 (EUbOPEN project). The IMI2 Joint Undertaking receives support from the European Union’s Horizon 2020 research and innovation programme, European Federation of Pharmaceutical Industries and Associations (EFPIA) companies, and associated partners KTH, OICR, Diamond and McGill. Biophysics, drug discovery, proteomics, and computing activities at Dundee were partially supported by Wellcome Trust strategic awards (100476/Z/12/Z, 094090/Z/10/Z, and 097945/C/11/Z, respectively) and the University of Dundee. A.D.C. was supported by a Horizon2020 Marie Skłodowska–Curie Actions Individual Fellowship (H2020-MSCA-IF-2020-101024945 DELETER).

### Authors’ contributions

A.D.C. and A.C. developed the concept of the study, S.J. performed initial E3 scoping to select DCAF proteins. A.D.C. designed and performed all experiments with input from A.C. A.D.C. wrote the initial manuscript, A.D.C. and A.C. edited the manuscript and all authors approved the final manuscript.

## Acknowledgements

We acknowledge the technical and research staff of the Ciulli Laboratories at CeTPD for the set-up and upkeep of protein expression and purification infrastructure. We thank Kwok-Ho Chan (Ciulli Lab) for preliminary work on DCAF expression, and Tasuku Ishida (Ciulli Lab) for sharing of insect cell reagents and resources; the MRC PPU Reagents and Services facility (MRC PPU, School of Life Sciences, University of Dundee, Scotland, mrcppureagents.dundee.ac.uk) for the reagents and services indicated in this publication, particularly Rachel Toth and Nicola Wood for cloning work; and Sharon Shepherd (BCDD, School of Life Sciences, University of Dundee) for advice and equipment access for insect cell expression. ChatGPT was used to improve readability of the manuscript. Figure 2 was created in BioRender https://BioRender.com/h4n1nof.

