## Supplementary Figures for "Scalable insect cell expression and purification screening applied to CRL4-DCAF substrate receptors"

**Supplementary Figures 1 to 9** – SDS-PAGE gels of nickel affinity purified DCAF constructs. The MRC PPU clone numbers, construct names and expected molecular weights are provided. Baculoviruses included in each sample lane are indicated by filled black dots. Individual DCAF protein constructs are divided into subsets of 4 lanes, samples in the first two lanes were infected with DCAF virus only, the 3rd lane was infected with DCAF virus and a virus encoding both DDB1 and DDA1, and the 4th lane was infected with DCAF virus and a virus encoding both DDB1$^{\Delta BPB}$ and DDA1. DDB1, DDB1$^{\Delta BPB}$ and DDA1 band positions and expected molecular weights are labelled on the right-hand side of the gels. Despite running at slightly different apparent molecular weights from expected weights, masses of DDB1, DDB1$^{\Delta BPB}$ and DDA1 produced from these baculoviruses were validated by intact mass spec to be of the expected molecular weights (data not shown).

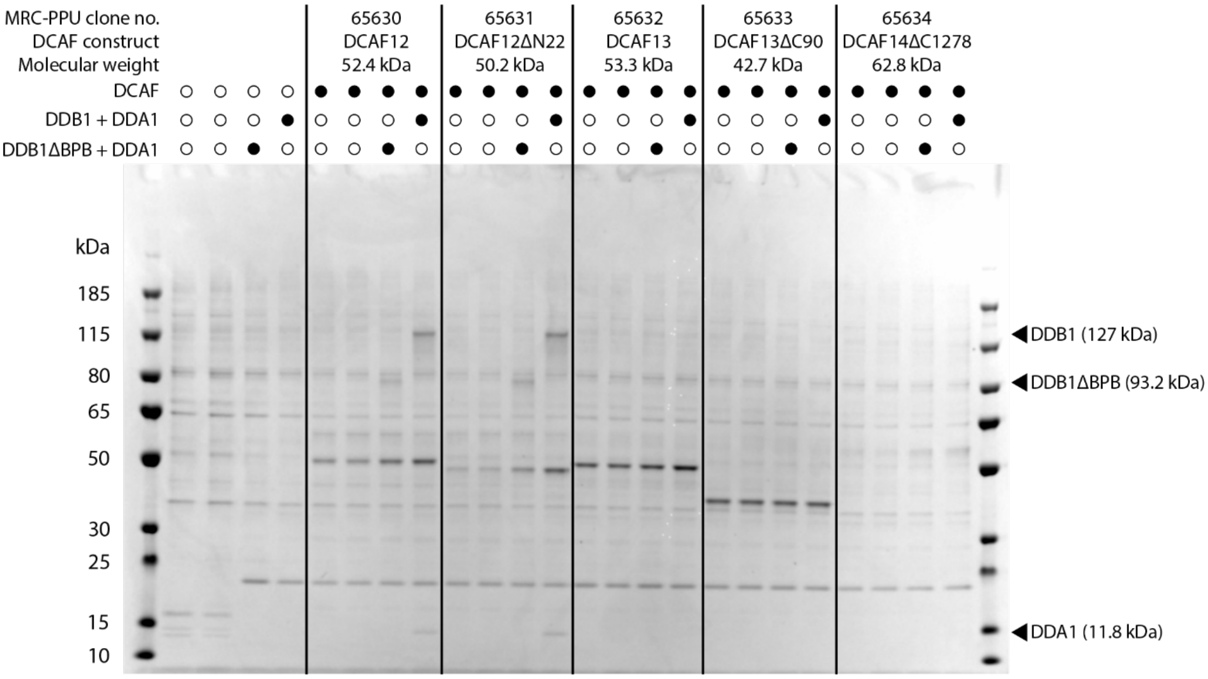


*Supplementary Figure 1.* SDS-PAGE gel of nickel affinity purified DCAF constructs DCAF12, DCAF12ΔN22, DCAF13, DCAF13ΔC90 and DCAF14ΔC1278.

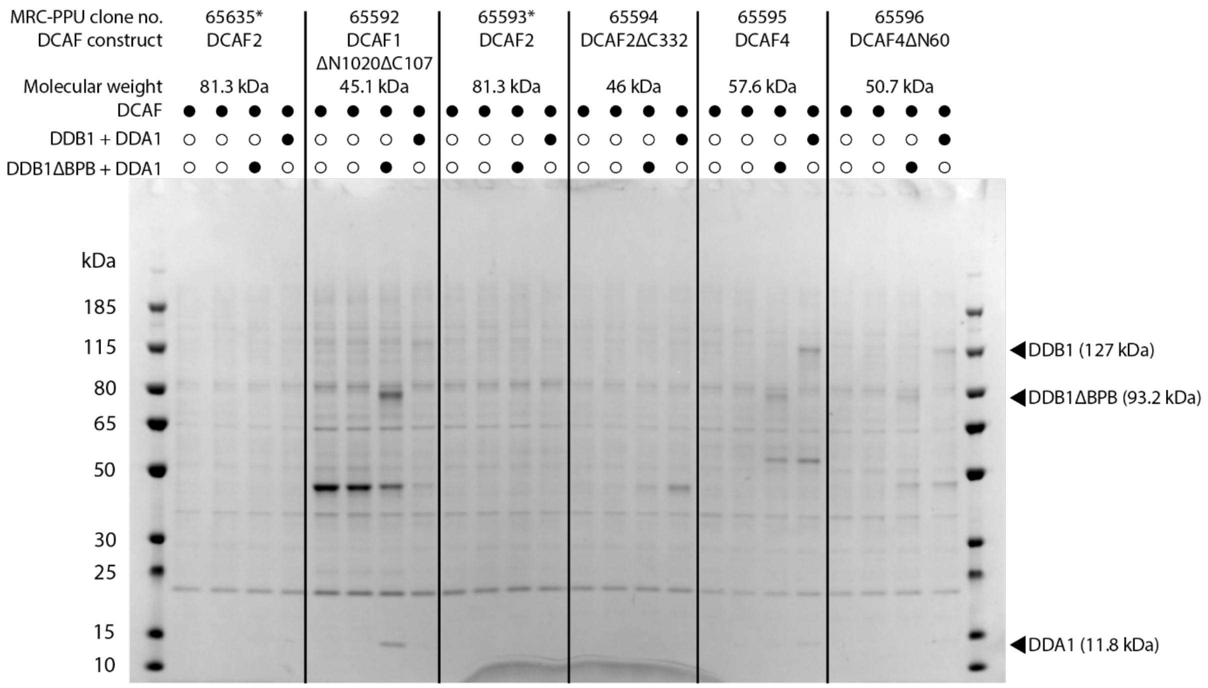


*Supplementary Figure 2.* SDS-PAGE gel of nickel affinity purified DCAF constructs DCAF2* (same construct ordered and run twice by accident), DCAF1ΔN1020ΔC107, DCAF2ΔC332, DCAF4 and DCAF4ΔN60.

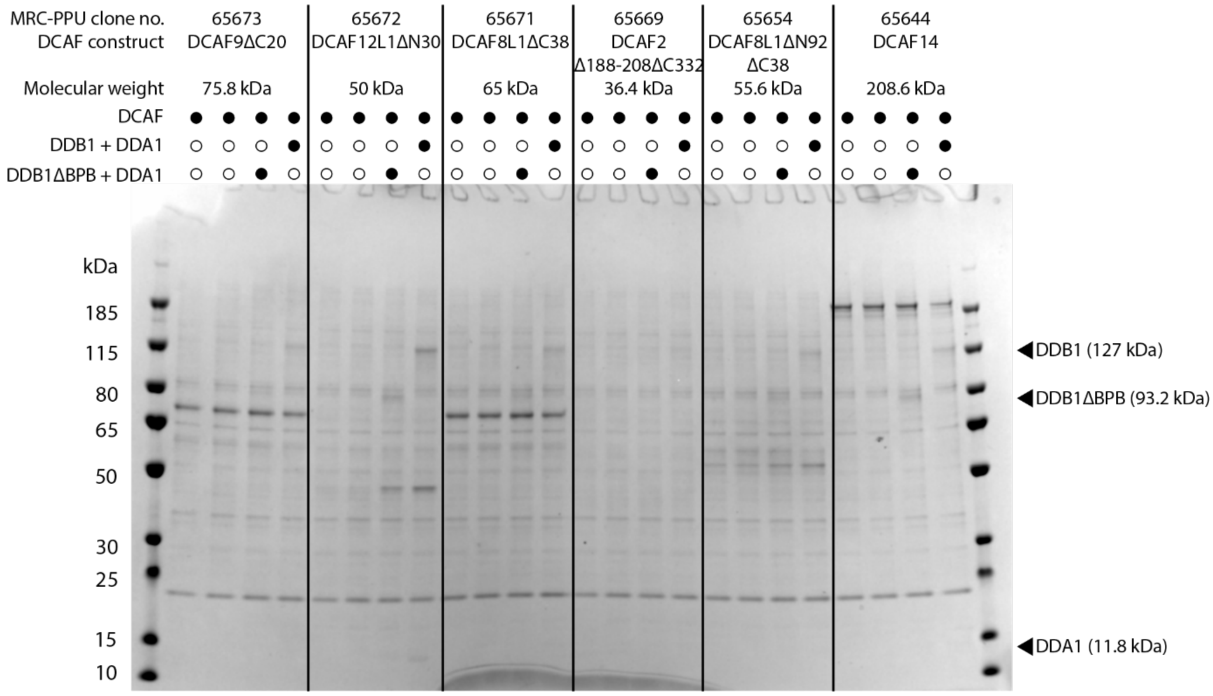


*Supplementary Figure 3.* SDS-PAGE gel of nickel affinity purified DCAF constructs DCAF9ΔC20, DCAF12L1ΔN30, DCAF8L1ΔC38, DCAF2Δ188-108ΔC332, DCAF8L1ΔN92ΔC38 and DCAF14.

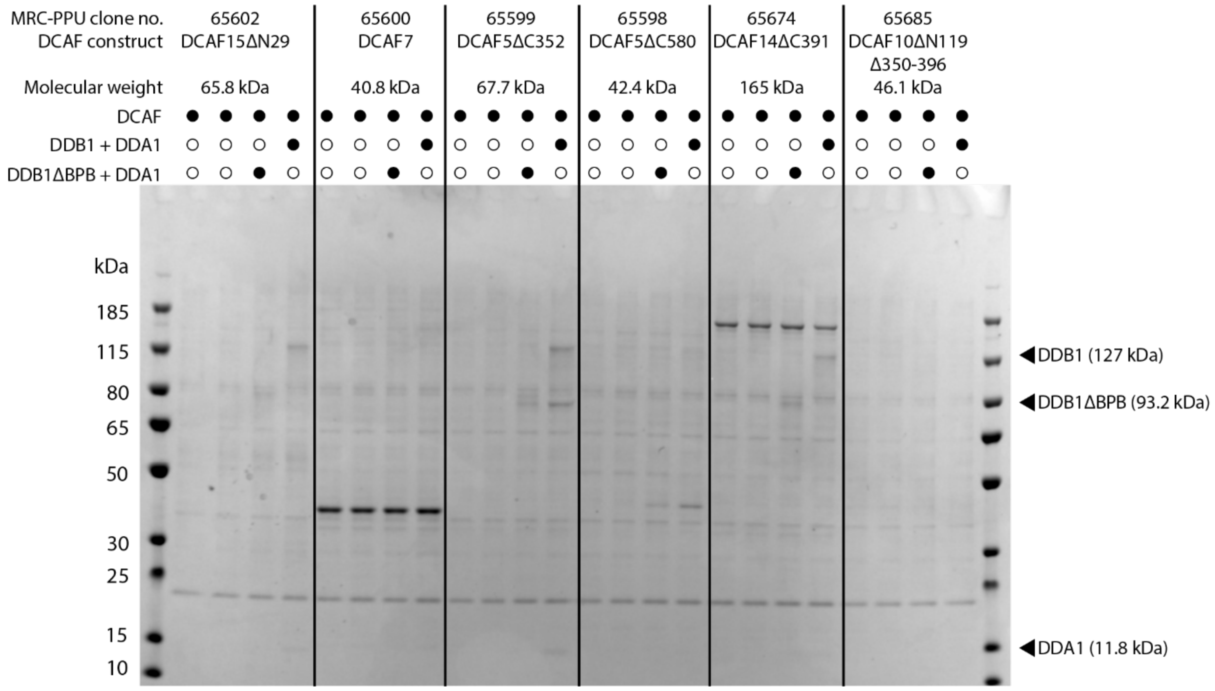


*Supplementary Figure 4.* SDS-PAGE gel of nickel affinity purified DCAF constructs DCAF15ΔN29, DCAF7, DCAF5ΔC352, DCAF5ΔC580, DCAF14ΔC391 and DCAF10ΔN119Δ350-396.

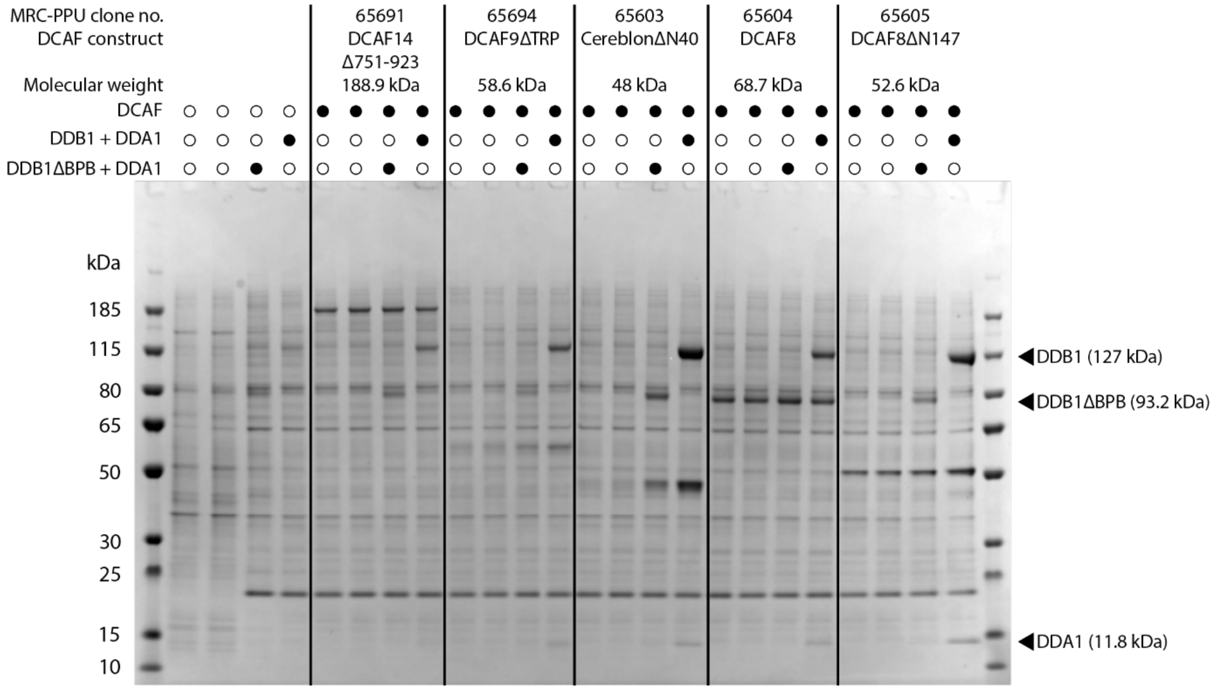


*Supplementary Figure 5.* SDS-PAGE gel of nickel affinity purified DCAF constructs DCAF14Δ751-923, DCAF9ΔTRP, CereblonΔN40, DCAF8 and DCAF8ΔN147.

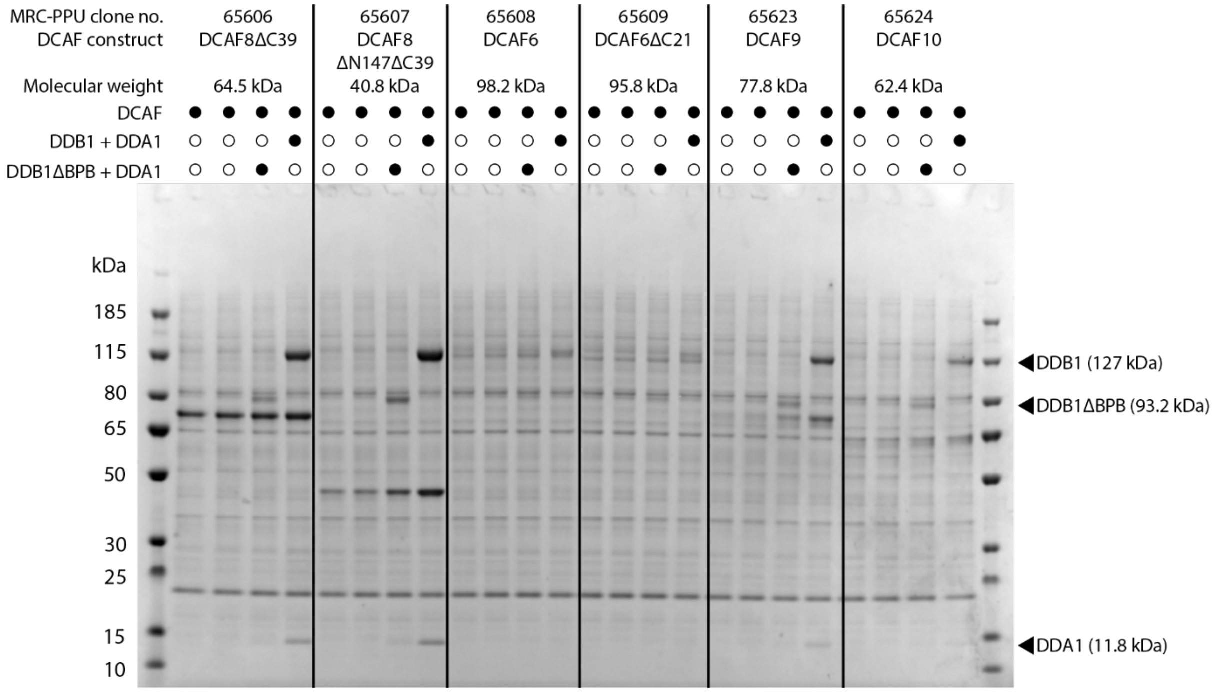


*Supplementary Figure 6.* SDS-PAGE gel of nickel affinity purified DCAF constructs DCAF8ΔC39, DCAF8ΔN147ΔC39, DCAF6, DCAF6ΔC21, DCAF9 and DCAF10.

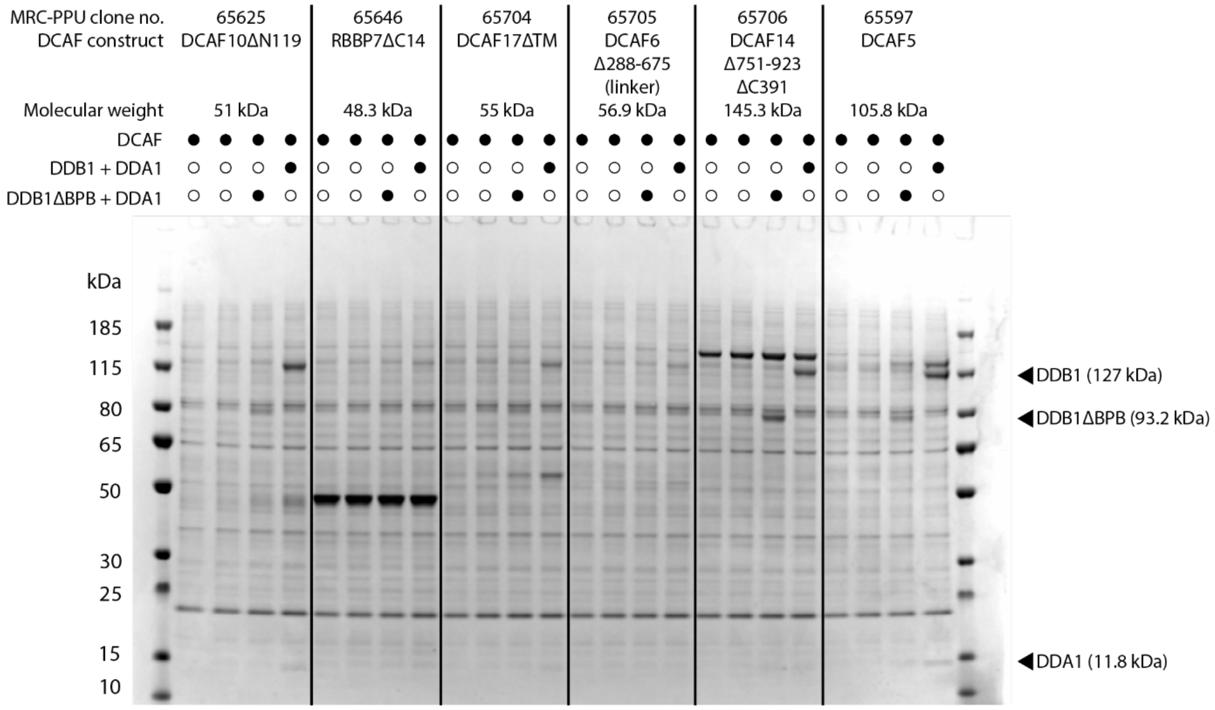


*Supplementary Figure 7.* SDS-PAGE gel of nickel affinity purified DCAF constructs DCAF10ΔN119, RBBP7ΔC14, DCAF17ΔTM, DCAF6Δ288-675(linker), DCAF14Δ751-923ΔC391 and DCAF5.

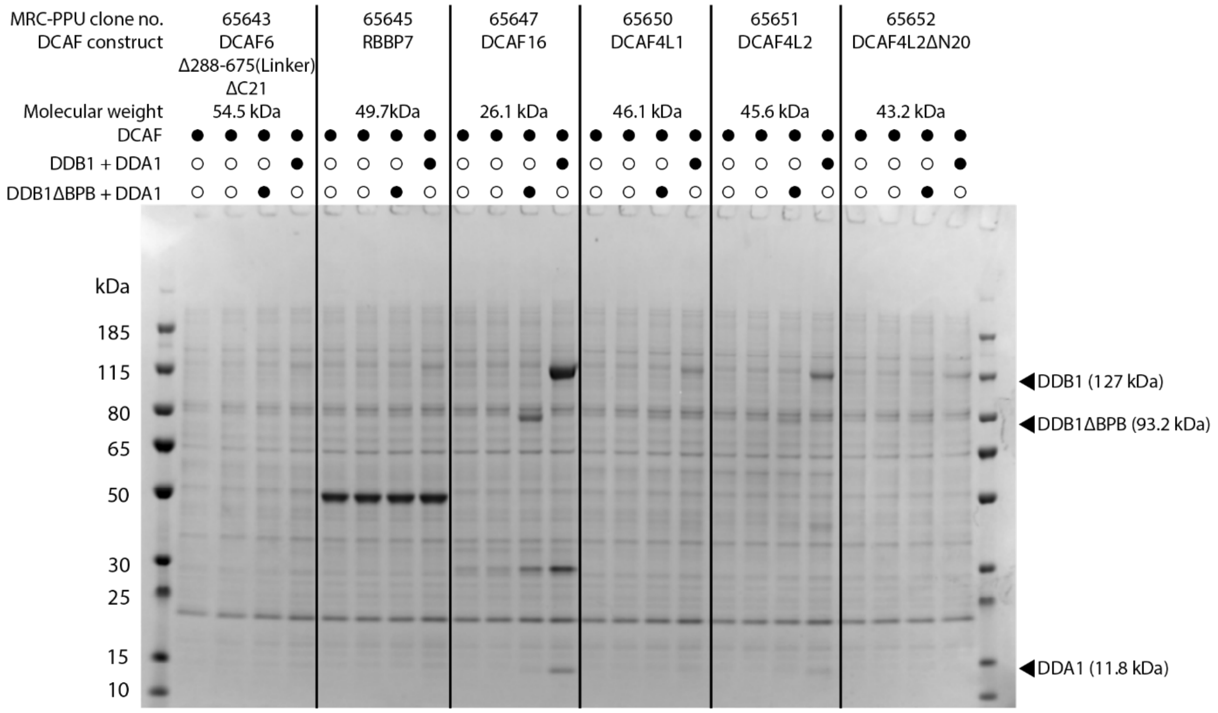


*Supplementary Figure 8.* SDS-PAGE gel of nickel affinity purified DCAF constructs DCAF6Δ288-675(linker)ΔC21, RBBP7, DCAF16, DCAF4L1, DCAF4L2 and DCAF4L2ΔN20.

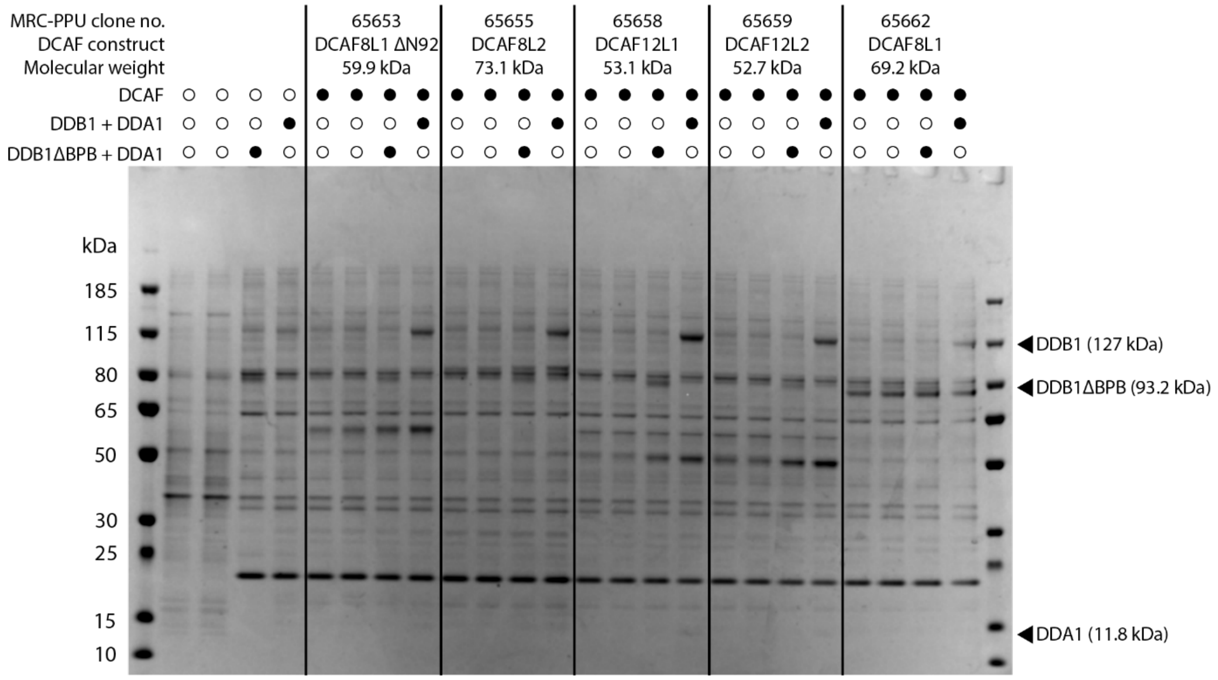


*Supplementary Figure 9.* SDS-PAGE gel of nickel affinity purified DCAF constructs DCAF8L1ΔN92, DCAF8L2, DCAF12L1, DCAF12L2 and DCAF8L1.

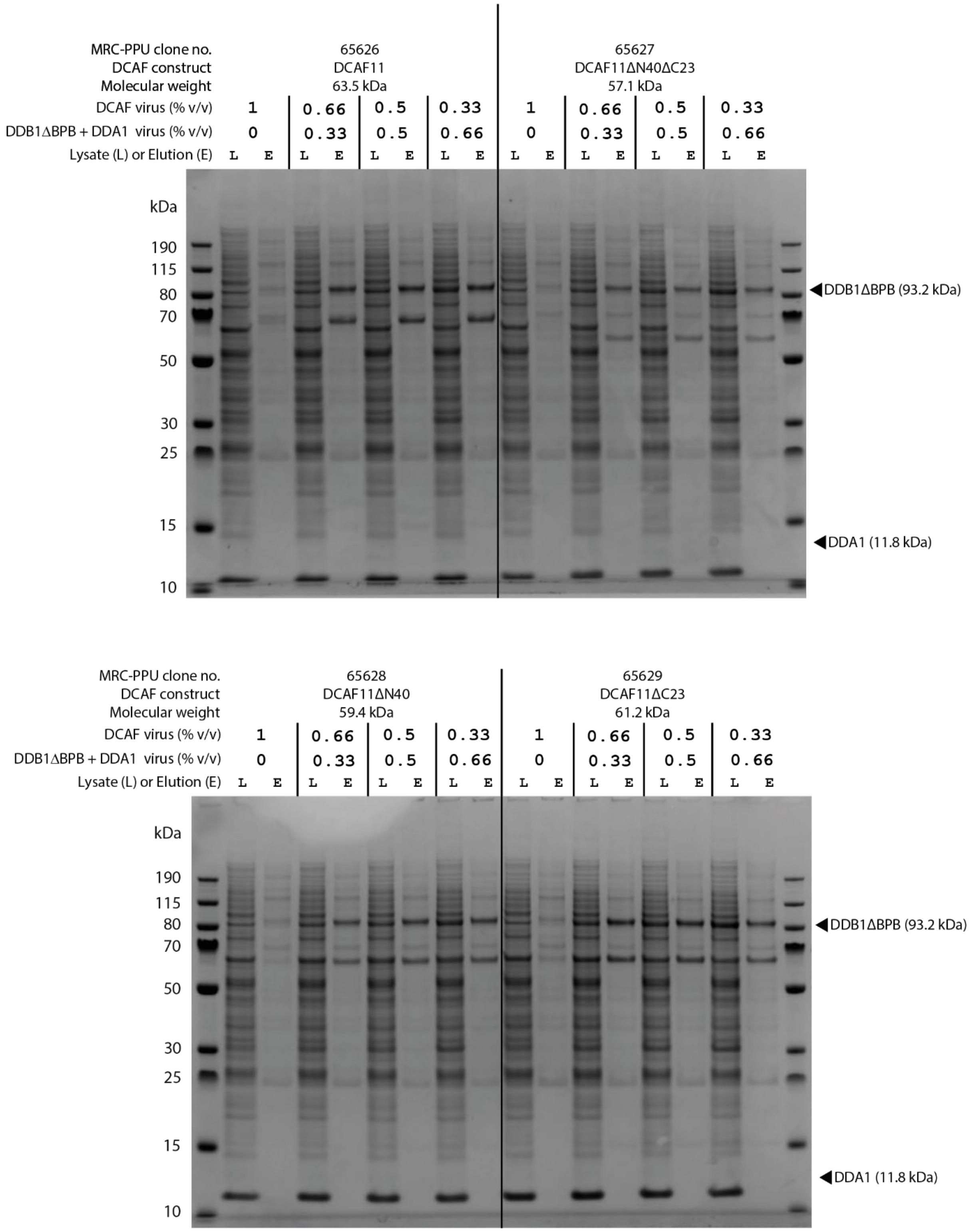


*Supplementary Figure 10.* SDS-PAGE gels (in this case run in MES instead of MOPS running buffer) of lysate and nickel affinity purified (elution) DCAF11 constructs. The MRC PPU clone numbers and expected molecular weights are provided. Viruses included in each sample lanes are indicated by a number that gives % viral volume (% v/v) added to the culture. Individual DCAF11 constructs are divided into subsets of 8 lanes, samples in the first two lanes were infected with only DCAF virus (1% v/v), and subsequent lanes were infected with different ratios of DCAF virus and virus expressing both DDB1ΔBPB and DDA1 with a total of 1% v/v. DDB1ΔBPB and DDA1 band positions and expected molecular weights are labelled on the right-hand side of the gel.
